# Assessment of Dried Blood Spots for DNA Methylation Profiling

**DOI:** 10.1101/546606

**Authors:** Rosie M. Walker, Louise MacGillivray, Sarah McCafferty, Nicola Wrobel, Lee Murphy, Shona M. Kerr, Stewart W. Morris, Archie Campbell, Andrew M. McIntosh, David J. Porteous, Kathryn L. Evans

**Affiliations:** Medical Genetics Section, Centre for Genomic and Experimental Medicine, Institute of Genetics and Molecular Medicine, University of Edinburgh, Edinburgh, EH4 2XU, UK; Centre for Cognitive Ageing and Cognitive Epidemiology, University of Edinburgh, Edinburgh, EH8 9JZ, UK; Edinburgh Clinical Research Facility, Western General Hospital, University of Edinburgh, Edinburgh, EH4 2XU, UK; MRC Human Genetics Unit, Institute of Genetics and Molecular Medicine, University of Edinburgh, Edinburgh, EH4 2XU, UK; Generation Scotland, Centre for Genomic and Experimental Medicine, Institute of Genetics and Molecular Medicine, University of Edinburgh, Edinburgh, EH4 2XU, UK; Division of Psychiatry, University of Edinburgh, Royal Edinburgh Hospital, Edinburgh, EH10 5HF, UK

**Author notes:** Equal last authors (David J. Porteous, Kathryn L. Evans).

## Abstract

**Background:** DNA methylation reflect health-related environmental exposures and genetic risk, providing insights into aetiological mechanisms and potentially predicting disease onset, progression and treatment response. An increasingly recognised need for large-scale, longitudinally-profiled samples collected world-wide has made the development of efficient and straightforward sample collection and storage procedures a pressing issue. An alternative to the low-temperature storage of EDTA tubes of venous blood samples, which are frequently the source of the DNA used in such studies, is to collect and store at room temperature blood samples using filter paper engineered for the purpose, such as Whatman FTA^®^ cards. Our goal was to determine whether DNA stored in this manner can be used to generate DNA methylation profiles comparable to those generated using blood samples frozen in EDTA tubes.

**Methods:** DNA methylation profiles were obtained from matched EDTA tube and Whatman FTA^®^ card whole-blood samples from 62 Generation Scotland: Scottish Family Health Study participants using the Infinium HumanMethylation450 BeadChip. Multiple quality control procedures were implemented, the relationship between the two sample types assessed, and EWASs performed for smoking status, age and the interaction between these variables and sample storage method. Results: Dried blood spot (DBS) DNA methylation profiles were of good quality and DNA methylation profiles from matched DBS and EDTA tube samples were highly correlated (mean r = 0.991) and could distinguish between participants. EWASs replicated established associations for smoking and age, with no evidence for moderation by storage method.

**Conclusions:** Our results support the use of Whatman FTA^®^ cards for collecting and storing blood samples for DNA methylation profiling. This approach is likely to be particularly beneficial for large-scale studies and those carried out in areas where freezer access is limited. Furthermore, our results will inform consideration of the use of newborn heel prick DBSs for research use.

## Introduction

Recent technological advances have facilitated the high-throughput genome-wide measurement of DNA methylation, yielding many studies reporting links between methylation and several conditions, diseases and phenotypes. The majority of studies have profiled DNA methylation in peripheral blood due to ease of sampling, which also renders it a useful tissue for identifying biomarkers. Blood samples for DNA methylation profiling are typically frozen prior to DNA extraction, with some samples being stored for years before use. As the sample numbers used for studies of DNA methylation, particularly epigenome-wide association studies (EWASs), continue to grow and the value of repeat sampling over time and exposures is recognised, so the logistical and financial issues surrounding whole blood sample collection and storage escalate. Moreover, population differences in DNA methylation^[1–4]^ highlight the need to profile DNA methylation around the world, including in low- and middle-income countries and rural settings, which might lack the required facilities for the low-temperature storage of EDTA blood collection tubes.

A potential alternative that would permit field collection and storage as well as postal collection is to collect finger or heel prick blood samples on filter paper^[5]^. As part of Generation Scotland: Scottish Family Health Study (GS:SFHS)^[6, 7]^, Whatman FTA^®^ cards were used to spot 100μl of the peripheral (venous) blood samples obtained at the baseline research clinic appointment, which took place between 2006 and 2011. These cards contain chemicals that lyse cell membranes and denature proteins, thus protecting DNA from degradation (https://www.sigmaaldrich.com/catalog/product/sigma/whawb120205?lang=en&region=GB). Samples collected on these cards can be stored at room temperature for several years^[8]^ prior to DNA extraction, thus avoiding the financial and logistical issues associated with freezer storage. Whatman FTA^®^ cards have been validated for genetic studies^[9–11]^; however, the effect of storing samples in this manner on the epigenome is less well-established. Previous studies provide support for the possibility of profiling DNA methylation in samples stored on Whatman FTA^®^ cards^[8, 12]^; however, these studies were limited by their focus on either a small number of individuals (n = 2)^[12]^ or a single gene of interest^[8]^ and a relatively short time interval between sample collection and DNA extraction (maximum of approximately two years). Moreover, to the best of our knowledge, no previous study has compared genome-wide DNA methylation profiles obtained from dried blood spots (DBSs) stored on Whatman FTA^®^ cards with those from DNA from matched whole blood samples frozen in EDTA tubes (henceforth referred to as “EDTA samples”).

The matched DBSs and EDTA samples obtained from GS:SFHS^[6, 7]^ have permitted us to carry out this comparison using samples from 62 participants. We show that it is technically feasible to measure DNA methylation from DBSs stored on Whatman FTA^®^ cards, that the methylation profiles of DBS samples are highly correlated with those from EDTA samples from the same individuals, and that EWASs using DBS samples replicate well-established differentially methylated loci associated with smoking and age. These findings indicate that this storage method represents a viable alternative to freezing blood samples for DNA methylation studies.

In addition to supporting the use of DBSs as a solution to the challenges posed by the need to collect and store large numbers of blood samples for DNA methylation studies in a wide-range of sociodemographic settings, our results also provide encouragement for the ability to profile DNA methylation in DBSs routinely collected on filter paper (commonly referred to as Guthrie cards) from new-born babies. The Guthrie cards are used clinically to screen for a prescribed set of metabolic and genetic disorders. These tests do not exhaust the sample and the remaining material is typically stored, often indefinitely. The ability to profile DNA methylation from Guthrie cards would present a number of important opportunities^[13, 14]^, including: addressing the issues of reverse causation and confounding, which plague EWASs; anchoring population studies of gene-environment interactions at the beginning of life; and permitting the development of predictors of future disease onset, thus facilitating early intervention.

In Denmark, research access to Guthrie cards is possible through the Danish Newborn Screening Biobank^[15]^ and in 2017 it was estimated that almost 40% of the 2.1 million samples stored in this biobank had contributed to published research^[16]^. In Scotland, however, a moratorium on research access to Guthrie cards was imposed in 2009 and remains in place pending a public consultation^[17]^. An important step in determining whether medical researchers will be permitted access to this resource (over 2.5 million cards collected since 1965, including 9,788 GS:SFHS participants with genome-wide genotype data and extensive phenotype data) will be assessments of the feasibility of obtaining biologically meaningful and clinically valuable data from archived DBSs. Although the Whatman FTA^®^ card is not identical to the card types used in the NHS Scotland new-born screening programme, some of the issues addressed in the present study, such as low DNA yield, a long period of storage, and the necessity of using a sample comprising heterogeneous cell types, are relevant to the potential use of Guthrie cards for research.

## Methods

### Cohort description

Genome-wide DNA methylation was profiled in whole blood samples from 62 GS:SFHS participants, chosen to have matched DNA samples from whole blood stored in EDTA tubes, DBSs stored on Whatman FTA^®^ cards and complete phenotype data (on sex, age and smoking status). GS:SFHS has been described in detail elsewhere^[6, 7]^; further information and information on research access can be found <u>here</u>. Studies arising from GS:SFHS can be found <u>here</u>. Briefly, GS:SFHS comprises ~24,000 individuals who were aged 18 years or over at the time of recruitment. At the time of blood sample collection, participants were deeply phenotyped for a range of demographic, health, and social variables. A favourable ethical opinion for GS:SFHS was obtained from the NHS Tayside Committee on Medical Research Ethics, on behalf of the National Health Service (reference: 05/S1401/89). GS:SFHS has Research Tissue Bank Status (reference: 15/ES/0040).

### Blood sample collection and DNA extraction

Each participant provided a blood sample (9ml), which was collected in an EDTA tube for DNA extraction, when attending the baseline GS:SFHS clinical appointment. This sample was stored at - 20°C for between 33 and 497 days. At this point, 100μl of the sample (approximately equivalent to two drops of blood) was blotted onto a Whatman FTA^®^ card (Sigma-Aldrich) and DNA was extracted from the remaining sample using the Nucleon BACC3 Genomic DNA Extraction Kit (Fisher Scientific)^[18]^, following the manufacturer’s instructions. The Whatman FTA^®^ cards were stored at room temperature for between 5 and 10 years until DNA extraction for this project. DNA was extracted from the entire blood spot using the QIAamp DNA Investigator Kit (Qiagen), following the manufacturer’s instructions. Sample storage and DNA extraction took place at the Edinburgh Clinical Research Facility at the University of Edinburgh (https://www.edinburghcrf.ed.ac.uk).

### Genome-wide methylation profiling

Whole blood genomic DNA (EDTA samples: 500ng; DBS samples: 212-500ng) was treated with sodium bisulphite using the EZ-96 DNA Methylation Kit (Zymo Research, Irvine, California), following the manufacturer’s instructions. DNA methylation was profiled using the Infinium HumanMethylation450 BeadChip (Illumina Inc.), according to the manufacturer’s protocol. Paired EDTA and DBS samples from a given participant were profiled on separate slides, matched for row position.

Raw intensity (.idat) files were read into R^[19]^ using the minfi package^[20]^ and quality control was carried out using shinyMethyl^[21]^ and functions within the wateRmelon package^[22]^. ShinyMethyl was used to create a “QC plot”, which plots the log median intensity of the unmethylated signal against the log median intensity of the methylated signal for each array, and density plots of the beta-values separated by probe type (type 1 or 2) and colour channel for the type 1 probes. These plots were assessed for outlier samples by visual inspection. The performance of each of the 14 types of control probe on the array was assessed by inspecting plots created by shinyMethyl showing the raw signal intensities for each array. Outliers were assessed by visual inspection. ShinyMethyl’s sex prediction plot was used to assess whether a participant’s predicted sex based on the difference between the median copy number intensity for the Y chromosome and the median copy number intensity for the X chromosome matched their self-reported sex. Next, the pfilter function in wateRmelon was used to exclude poor-performing samples and probes. Samples with ≥ 1% sites with a detection p-value of > 0.05 were excluded. Probes were excluded from the dataset if: (i) ≥5 samples had a beadcount of < 3; or (ii) ≥ 0.5% samples had a detection p-value of > 0.05. Probes that had been predicted to cross-hybridise were removed using the rmSNPandCH function in the R package DMRcate^[23]^, which removes cross-reactive probes identified by Chen et al.^[24]^. Probes on the X and Y chromosomes and those that target loci affected by a SNP at the CpG site or at the site of single base extension (in the case of Type 1 probes) were removed prior to carrying out EWASs.

The data from all samples were normalised together using the dasen method in wateRmelon. Dasen adjusts the background difference between Type I and Type II assays (by adding the offset between Type I and II probe intensities to Type I intensities) and then performs between-array quantile normalisation for the methylated and unmethylated signal intensities separately (Type I and Type II assays normalised separately).

### Assessment of the relationship between detection p-value and DBS sample characteristics

A paired t-test was performed to determine whether there was a difference in the total number of sites per sample with a poor detection p-value (detection p > 0.05) between the DBS and EDTA samples, with p ≤ 0.05 deemed to be significant. The relationships between the total number of sites for each sample with a detection p-value greater than 0.05 and (i) sample storage time and (ii) quantity of DNA hybridised to the array were assessed by linear modelling, with p ≤ 0.05 deemed to be significant.

### Estimation of whole blood cellular composition

Estimated cell counts for B-lymphocytes, granulocytes, monocytes, natural killer cells, CD4+ T-lymphocytes and CD8+ T-lymphocytes were obtained using the estimateCellCounts function in minfi. This function implements Jaffe and Irizarry’s^[25]^ modification of Houseman’s^[26]^ algorithm. Paired t-tests were performed to assess whether blood storage method (EDTA tube vs. Whatman FTA^®^ card) affected cellular composition. Correction for multiple testing was implemented using the Benjamini-Hochberg false discovery rate (FDR), with q-values of ≤ 0.05 deemed to be significant.

### Assessment of the correlation between methylation profiles obtained from EDTA tube and DBS paired samples and unsupervised hierarchical clustering

Following the method described by Hannon et al. (2015)^[27]^, a set of probes deemed to be variable in our sample was identified. The pair-wise correlation between all samples was assessed by calculating Pearson correlation coefficients. Unsupervised hierarchical clustering plots were generated using the R package lumi^[28]^ using the same set of variable probes as used for the correlation analysis.

### Identification of differentially methylated positions associated with smoking or age and the interaction with storage method

EWASs were carried out (i) to identify loci where DNA methylation is associated with smoking or age in the EDTA or DBS samples and (ii) to establish whether these associations were moderated by storage method. EWASs for smoking and age were implemented in the R package limma^[29]^ by fitting linear models in which DNA methylation (beta-value) was the outcome variable and age or smoking (a categorical variable with the following levels: current smoker, former smoker who gave up under 12 months ago, former smoker who gave up 12 months or longer ago, or never smoked) was the predictor of interest. When smoking status was the predictor of interest, the following covariates were included in the model: age, sex, slide ID, row number and estimated proportions of B-lymphocytes, granulocytes, natural killer cells, CD4+ T-lymphocytes and CD8+ T-lymphocytes. The same covariates were included for the EWAS of age with the addition of “pack years”, a measure that indicates an individual’s lifetime exposure to tobacco. Pack years were calculated by multiplying the years an individual had smoked for by the maximum number of packs of cigarettes they ever smoked per day (a pack = 20 cigarettes). A conversion was used for cigars (a cigar = four cigarettes) and rolling tobacco (a 25g pack = 50 cigarettes). EWASs to assess the interaction between smoking status or age and storage method (EDTA tube or Whatman FTA^®^ card) were carried out using linear mixed effects models implemented in the R package nlme^[30]^. The fixed effects were the same as for the main effects EWASs, with the addition of a variable representing storage method, and each participant was modelled with a random intercept. The threshold for significance was defined as p ≤ 3.6 x 10^-8^, as recommended by Saffari et al. (2018)^[31]^.

## Results

### Extraction of DNA from DBSs stored on Whatman FTA^®^ cards

The concentration of the DNA extracted from the DBSs ranged from 3.8 ng/μl to 145.8 ng/μl, with a mean yield of 18.0 ng/μl, in a total volume of 50 μl. The recommended minimum input for the Infinium HumanMethylation450 BeadChip, 500 ng DNA, was available for 43 of the 62 samples, with all but two samples having at least 250 ng DNA as input. The lowest array input was 212 ng DNA.

### Quality control of DNA methylation arrays

Initial assessment of array performance using the R package shinyMethyl indicated that all arrays had performed well. Neither the QC plot (Figure 1) nor the density plots of M-values and beta-values (Figure 2) produced using shinyMethyl indicated any obvious outliers. Both plot types did, however, indicate a slight shift between the EDTA and DBS samples. From the QC plot it can be seen that there was a tendency for the DBS samples to have slightly lower methylated signal intensity (y axis) and a higher unmethylated signal intensity (x axis). From the density plots, a slight shift in distributions, which was most evident at the extremes of the beta-value range (i.e. 0 and 1), could be seen for type 1 probes in both the green (Figure 2A) and red (Figure 2B) colour channels and type 2 probes (Figure 2C). This shift was most obvious for type 2 probes at the methylated end of the beta-value range, such that fewer sites in DBS samples showed the highest levels of DNA methylation. These differences in overall signal intensity were not, however, reflected by differences in the performance of the control probes. Plots of the output of these probes supported the conclusion that all arrays had performed well, with no obvious outliers (Figure 3). Importantly, the bisulphite conversion control probes did not indicate a difference in bisulphite conversion efficiency between EDTA and DBS samples and the non-polymorphic control probes, which measure overall assay performance, did not appear to differ according to sample storage method.

**Figure 1.**
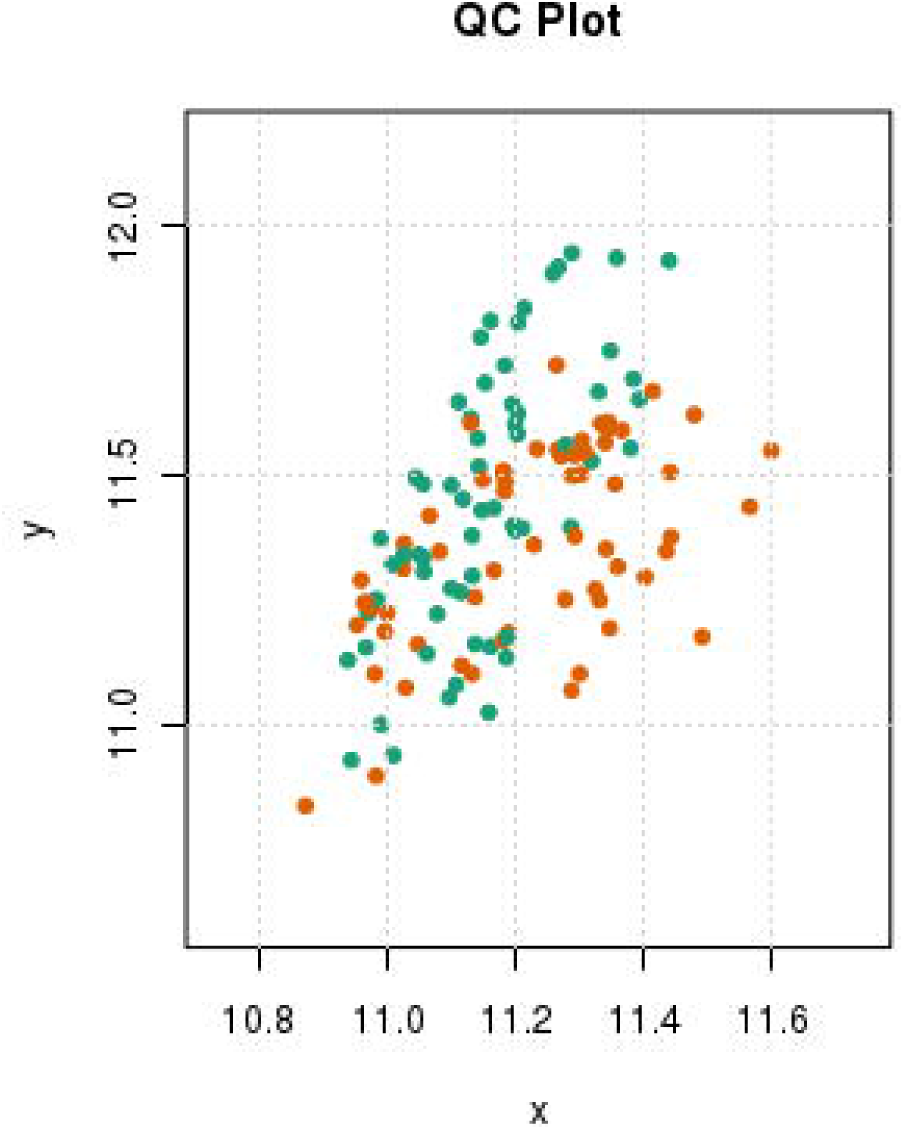
Quality control plot showing median methylated and unmethylated raw signal intensities. For each sample, the log median methylated signal intensity (y-axis) is plotted against the log median unmethylated signal intensity (x-axis). Each dot represents one sample with EDTA samples being represented in green and DBS samples in orange.

**Figure 2.**
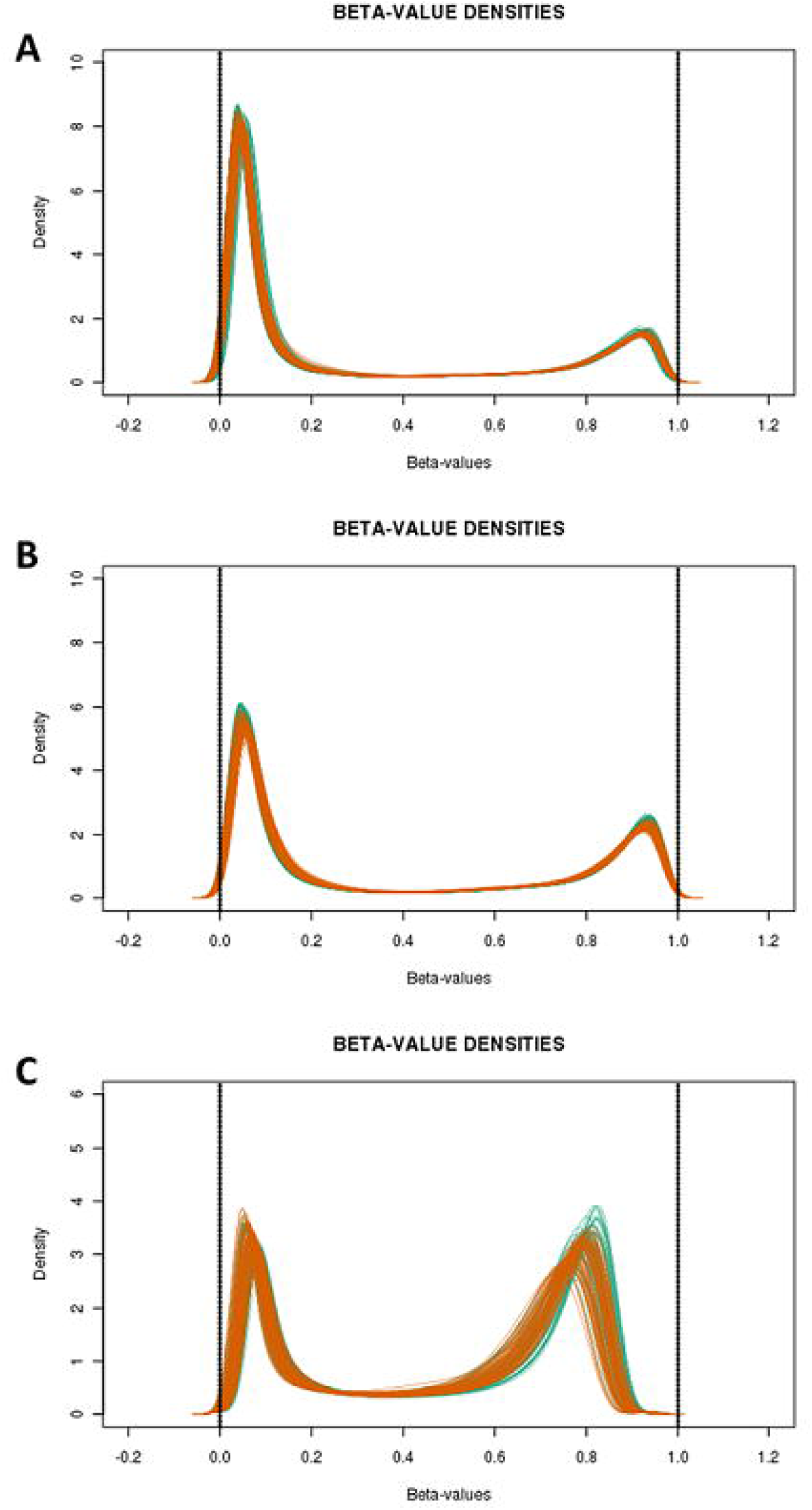
Density plots of the raw beta-values separated by probe type and methylation measurement unit. Density plots are shown for the type 1 (green colour channel (A); red colour channel (B)) and type 2 (C) probes. Each sample is represented by a separate line with EDTA samples indicated in green and DBS samples in orange.

**Figure 3.**
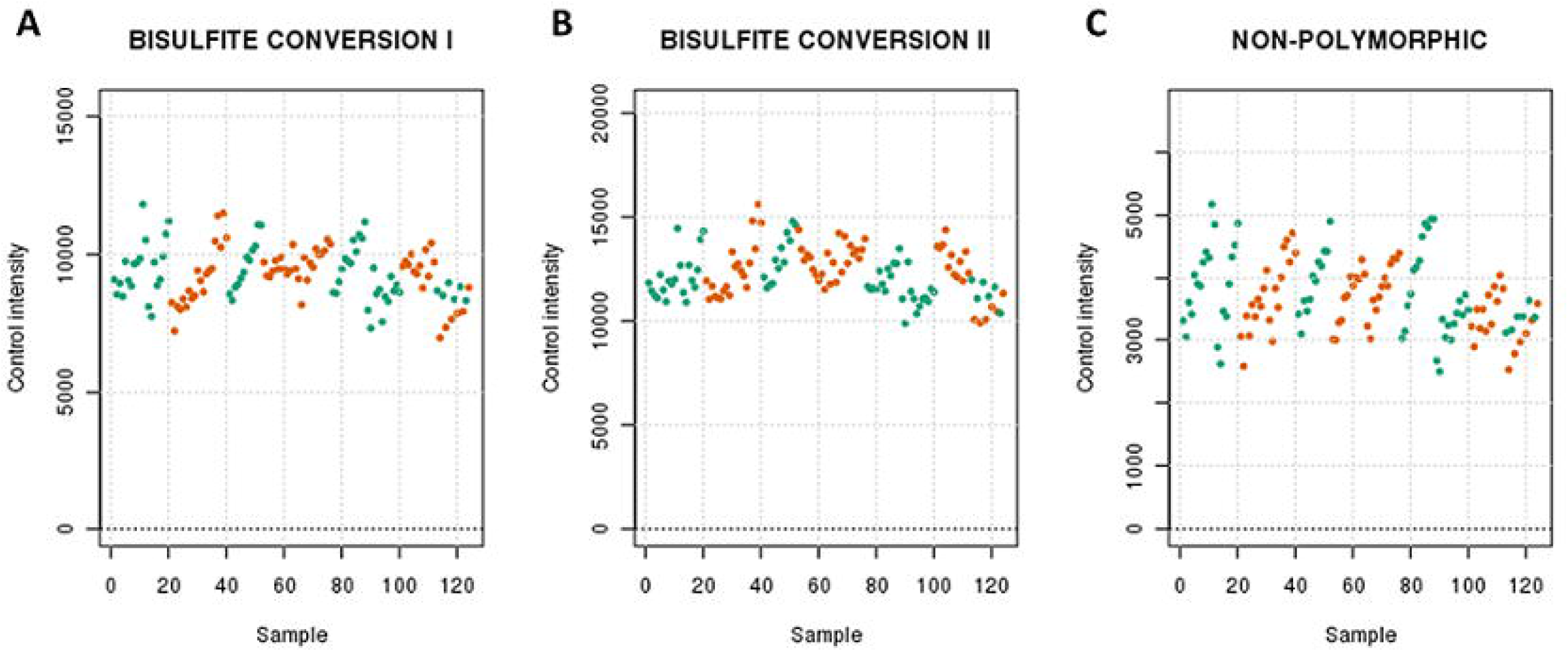
Control probe plots for the bisulphite conversion (I and II) and non-polymorphic probe control probes. The mean raw signal intensity for the bisulphite conversion (I and II) (A and B) and non-polymorphic (C) control probes are plotted for each array. EDTA samples are indicated in green and DBS samples in orange.

In keeping with these observations, no samples were excluded based on the proportion of poorly detected sites. There was, however, a significant difference between the DBS and EDTA blood samples in the number of sites per sample with a poor detection p-value (p = 0.0177), such that the DBS samples had fewer sites per sample with a poor detection p-value than the EDTA samples (mean difference = −63.7 sites/sample). For the DBS samples, the total number of sites per sample with a poor detection p-value was not related to storage time (p = 0.498) or the amount of DNA hybridised to the array (p = 0.220).

Two hundred and ninety-two sites were excluded as their beadcount was lower than three in at least five percent of samples and 2,280 sites were excluded due to having at least one percent of samples with a detection p-value greater than 0.05. Following the exclusion of probes that have been predicted to cross-hybridise, 452,278 probes were retained in the dataset, with this number being reduced to 436,309 after excluding probes on the X and Y chromosomes and those affected by a SNP at the CpG site or the site of single base extension (for type 1 probes).

### Participant and sample characteristics

The mean age of the participants was 31.8 years (range = 18.1-42.8 years; standard deviation = 7.2 years) and 62.9% of the sample were female. Fourteen participants were current smokers and 32 had never smoked. The mean storage time for the DBS samples was 7.78 years (range = 5.79 – 9.96 years).

### Assessment of the effect of storage method on whole blood cellular composition

Sample storage method (EDTA vs. DBS) was not observed to be associated with the proportion of any of the six estimated blood cell types (min. q = 0.103; Table 1).

**Table 1.** Assessment of the relationship between blood cell type proportions and sample storage method. Six blood cell types were estimated using the estimateCellCounts function in the R package minfi and associations with method of blood sample storage (EDTA tube vs. Whatman FTA card) were assessed by paired t-tests. A Benjamini-Hochberg FDR adjustment (q-value) was applied to account for the number of tests carried out and statistical significance was defined as q ≤ 0.05.

| Cell type | t-statistic | *p*-value | *q*-value |
| --- | --- | --- | --- |
| B-cells | -0.63 | 0.528 | 0.736 |
| CD4+ T-cells | -1.56 | 0.124 | 0.373 |
| CD8+ T-cells | 2.45 | 0.0171 | 0.103 |
| Granulocytes | 0.34 | 0.736 | 0.736 |
| Monocytes | 0.40 | 0.687 | 0.736 |
| Natural Killer cells | -1.19 | 0.240 | 0.480 |

### Assessment of the comparability of DNA methylation profiles obtained from DBS and EDTA tube-stored whole blood samples

Using a subset of probes defined as showing variation in our sample (n = 190,044), Pearson correlation coefficients were calculated between all pairs of samples. For all participants, the highest correlation coefficient was observed between the two blood sample types (EDTA and DBS) from that participant (mean r = 0.991, range = 0.976 - 0.995). For comparison, the pairwise correlation coefficients between pairs of samples from different participants (across both storage methods) had a mean of r = 0.953 and ranged from 0.904 – 0.976. Consistent with these observations, unsupervised hierarchical clustering performed using the same variable set of probes indicated a high level of similarity between the DBS and EDTA samples from the same participant, with methylation differences being sufficient to distinguish between participants (Figure 4).

**Figure 4.**
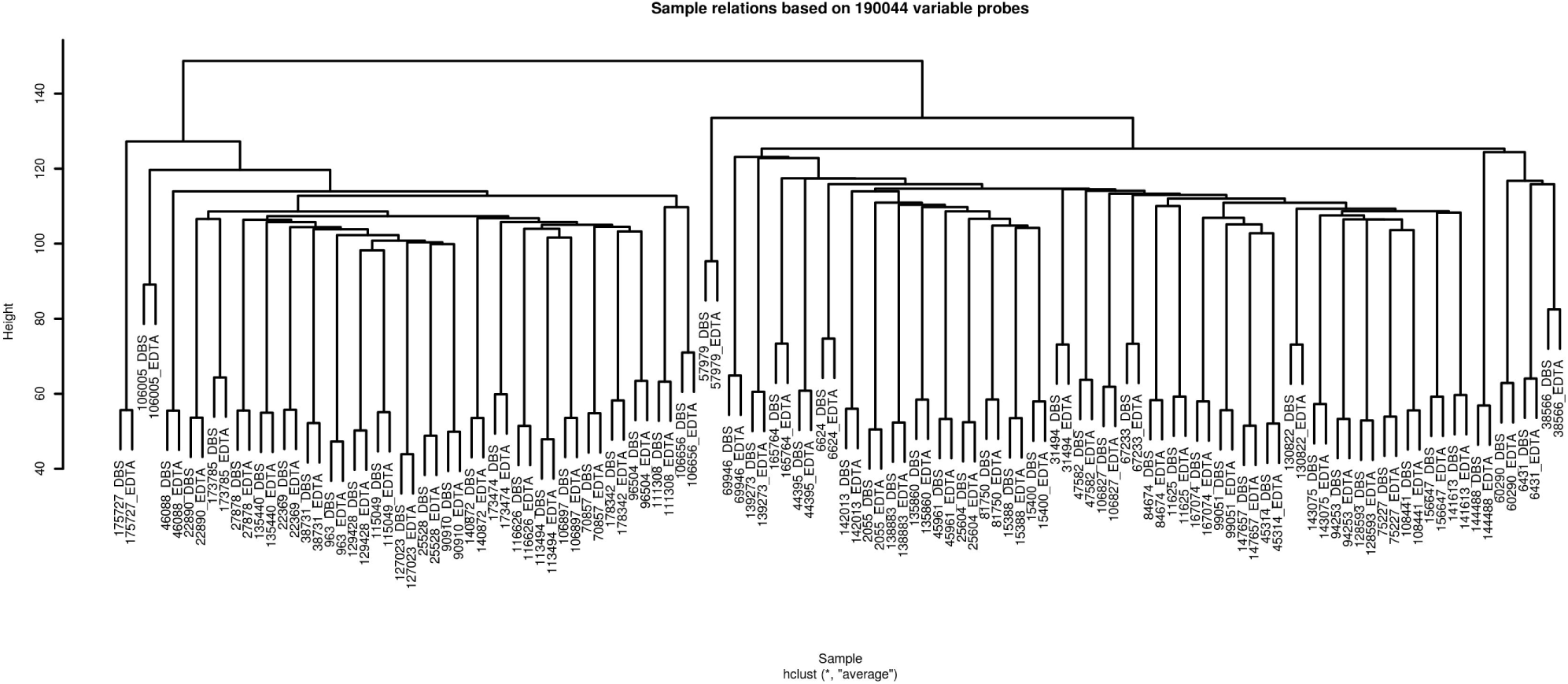
Unsupervised hierarchical clustering of methylation levels (beta-values) at variable probes. A subset of 190,044 probes defined as variable using an approach described by Hannon et al. (2015)^[27]^ were used to perform unsupervised hierarchical clustering of the 124 samples. Each sample is labelled using the participants study identifier and a suffix indicating whether the sample was from DNA from whole blood stored as a DBS or in an EDTA tube.

### Assessment of the relationship between DNA methylation and Whatman FTA^®^ card storage time

An EWAS was performed to establish whether the length of time that a sample is stored on a Whatman FTA^®^ card is associated with altered DNA methylation. This did not reveal any significant differences (min. q = 0.999).

### Assessment of the effect of sample storage method on the relationship between DNA methylation and age and smoking

Age and smoking have been robustly associated with altered DNA methylation by several previous studies. EWASs were carried out to (i) identify age- and smoking-associated differentially methylated positions (DMPs) in the EDTA and DBS samples when considered separately and (ii) assess whether sample storage method exerted an effect on these associations. EWASs for age and smoking status identified significant DMPs in both the EDTA and the DBS samples (Tables 2-5), while no significant associations were identified between DNA methylation and the interaction between storage method and age or smoking (min q = 0.933).

**Table 2.**
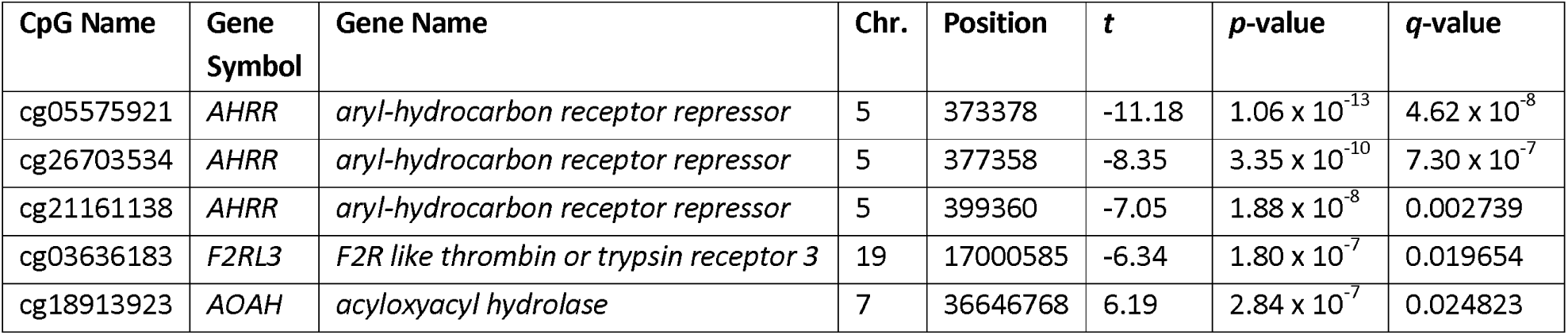
Significant results from an EWAS comparing current and never smokers carried out using DBS samples. For loci attaining a Benjamini-Hochberg q-value of ≤ 0.05, the CpG name, the symbol and name of the gene that the CpG is located in (or NA if the CpG is not located in a gene), genomic coordinate of the CpG (GRCh37), t-statistic, uncorrected p-value and q-value are shown. Results are ranked by p-value.

**Table 3.**
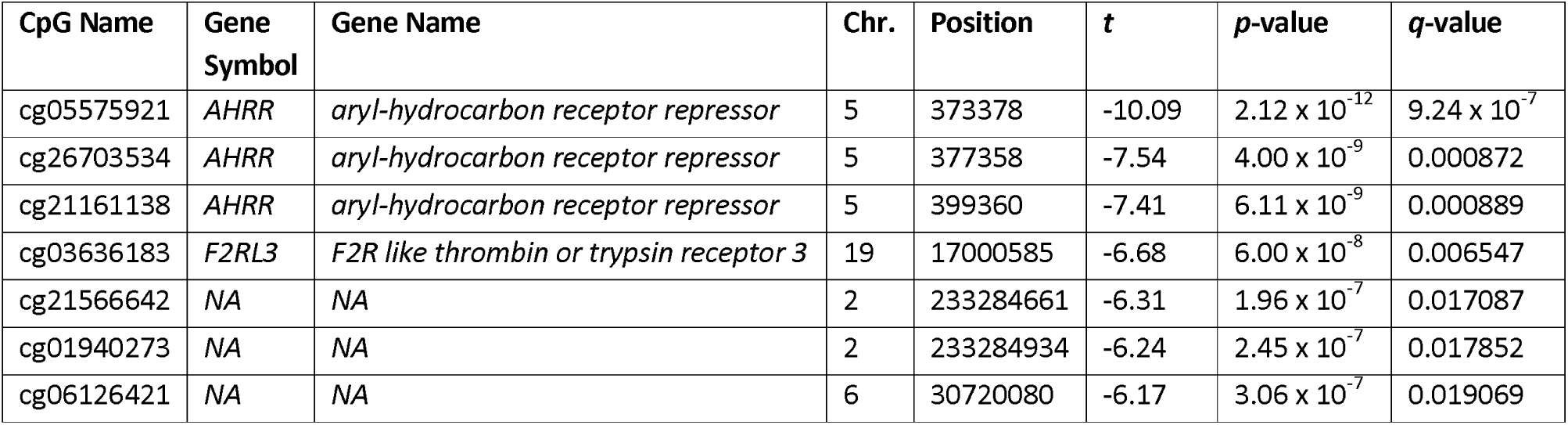
Significant results from an EWAS comparing current and never smokers carried out using EDTA samples. For loci attaining a Benjamini-Hochberg q-value of ≤ 0.05, the CpG name, the symbol and name of the gene that the CpG is located in (or NA if the CpG is not located in a gene), genomic coordinate of the CpG (GRCh37), t-statistic, uncorrected p-value and q-value are shown. Results are ranked by p-value.

| CpG Name | Gene Symbol | Gene Name | Chr. | Position | $t$ | $p$-value | $q$-value |
| --- | --- | --- | --- | --- | --- | --- | --- |
| cg05575921 | AHRR | aryl-hydrocarbon receptor repressor | 5 | 373378 | -10.09 | $2.12 \times 10^{-12}$ | $9.24 \times 10^{-7}$ |
| cg26703534 | AHRR | aryl-hydrocarbon receptor repressor | 5 | 377358 | -7.54 | $4.00 \times 10^{-9}$ | 0.000872 |
| cg21161138 | AHRR | aryl-hydrocarbon receptor repressor | 5 | 399360 | -7.41 | $6.11 \times 10^{-9}$ | 0.000889 |
| cg03636183 | F2RL3 | F2R like thrombin or trypsin receptor 3 | 19 | 17000585 | -6.68 | $6.00 \times 10^{-8}$ | 0.006547 |
| cg21566642 | NA | NA | 2 | 233284661 | -6.31 | $1.96 \times 10^{-7}$ | 0.017087 |
| cg01940273 | NA | NA | 2 | 233284934 | -6.24 | $2.45 \times 10^{-7}$ | 0.017852 |
| cg06126421 | NA | NA | 6 | 30720080 | -6.17 | $3.06 \times 10^{-7}$ | 0.019069 |

**Table 4.** The top 10 significant results from an EWAS of age carried out using DBS samples. For the first 10 loci attaining a Benjamini-Hochberg q-value of ≤ 0.05, the CpG name, the symbol and name of the gene that the CpG is located in (or NA if the CpG is not located in a gene), genomic coordinate of the CpG (GRCh37), t-statistic, uncorrected p-value and q-value are shown. Results are ranked by p-value.

| CpG Name | Gene Symbol | Gene Name | Chr. | Position | $t$ | $p$-value | $q$-value |
| --- | --- | --- | --- | --- | --- | --- | --- |
| cg16867657 | ELOVL2 | ELOVL fatty acid elongase 2 | 6 | 11044877 | 16.04 | $1.85 \times 10^{-18}$ | $8.09 \times 10^{-13}$ |
| cg10501210 | NA | NA | 1 | 207997020 | -9.39 | $2.01 \times 10^{-11}$ | $3.20 \times 10^{-6}$ |
| cg07553761 | TRIM59 | tripartite motif containing 59 | 3 | 160167977 | 9.36 | $2.20 \times 10^{-11}$ | $3.20 \times 10^{-6}$ |
| cg24724428 | ELOVL2 | ELOVL fatty acid elongase 2 | 6 | 11044888 | 8.97 | $6.65 \times 10^{-11}$ | $7.26 \times 10^{-6}$ |
| cg06639320 | FHL2 | four and a half LIM domains 2 | 2 | 106015739 | 8.75 | $1.26 \times 10^{-10}$ | $1.10 \times 10^{-5}$ |
| cg22454769 | FHL2 | four and a half LIM domains 2 | 2 | 106015767 | 8.58 | $2.07 \times 10^{-10}$ | $1.51 \times 10^{-5}$ |
| cg23606718 | AMER3 | APC membrane recruitment protein 3 | 2 | 131513927 | 7.71 | $2.86 \times 10^{-9}$ | 0.000178 |
| cg21572722 | ELOVL2 | ELOVL fatty acid elongase 2 | 6 | 11044894 | 7.41 | $7.11 \times 10^{-9}$ | 0.000388 |
| cg12934382 | GRM2 | glutamate metabotropic receptor 2 | 3 | 51741135 | 7.27 | $1.09 \times 10^{-8}$ | 0.000527 |
| cg08957484 | CCNI2 | cyclin I family member 2 | 5 | 132083532 | 7.20 | $1.36 \times 10^{-8}$ | 0.000592 |

**Table 5.** The top 10 significant results from an EWAS of age carried out using EDTA samples. For loci the first 10 loci attaining a Benjamini-Hochberg q-value of ≤ 0.05, the CpG name, the symbol and name of the gene that the CpG is located in (or NA if the CpG is not located in a gene), genomic coordinate of the CpG (GRCh37), t-statistic, uncorrected p-value and q-value are shown. Results are ranked by p-value.

| CpG Name | Gene Symbol | Gene Name | Chr. | Position | $t$ | $p$-value | $q$-value |
| --- | --- | --- | --- | --- | --- | --- | --- |
| cg16867657 | ELOVL2 | ELOVL fatty acid elongase 2 | 6 | 11044877 | 11.61 | $4.88 \times 10^{-14}$ | $2.13 \times 10^{-8}$ |
| cg22454769 | FHL2 | four and a half LIM domains 2 | 2 | 106015767 | 7.94 | $1.41 \times 10^{-9}$ | 0.000223 |
| cg10501210 | NA | NA | 1 | 207997020 | -7.91 | $1.53 \times 10^{-9}$ | 0.000223 |
| cg07547549 | SLC12A5 | solute carrier family 12 member 5 | 20 | 44658225 | 7.48 | $5.71 \times 10^{-9}$ | 0.000623 |
| cg16419235 | PENK | proenkephalin | 8 | 57360613 | 7.34 | $8.83 \times 10^{-9}$ | 0.000738 |
| cg06639320 | FHL2 | four and a half LIM domains 2 | 2 | 106015739 | 7.29 | $1.01 \times 10^{-8}$ | 0.000738 |
| cg21572722 | ELOVL2 | ELOVL fatty acid elongase 2 | 6 | 11044894 | 6.68 | $6.78 \times 10^{-8}$ | 0.004179 |
| cg23606718 | AMER3 | APC membrane recruitment protein 3 | 2 | 131513927 | 6.60 | $8.74 \times 10^{-8}$ | 0.004179 |
| cg17737621 | NA | NA | 20 | 21372480 | 6.60 | $8.80 \times 10^{-8}$ | 0.004179 |
| cg23500537 | NA | NA | 5 | 140419819 | 6.57 | $9.58 \times 10^{-8}$ | 0.004179 |

## Discussion

In agreement with previous studies, we have demonstrated that it is possible to obtain high quality DNA methylation profiles from archived DBSs stored on Whatman FTA^®^ cards and, for the first time, shown that these profiles are highly comparable to those obtained from matched whole blood DNA obtained from samples stored in EDTA tubes. Our findings support the use of Whatman FTA^®^ cards for the routine collection of blood samples for studies of DNA methylation. There are potentially substantial benefits to this approach, particularly for large-scale studies and those carried out in rural or remote settings and/or in low- and middle-income countries. The blood collection by finger prick protocol is comparable to that used for blood glucose monitoring in diabetes and could be self-administered by study participants.

DNA methylation profiles obtained from the DBSs appeared to be of good quality: on almost all comparison metrics, the performance of the DBS samples could not be distinguished from the performance of the EDTA tube samples. There was a tendency for DBS samples to have higher methylated signal intensity and lower unmethylated signal intensity than the EDTA samples, but this did not result in a difference in the performance of the control probes, or in the number of sites with a poor detection p-value. Encouragingly, the overall rate of site detection was slightly better in the DBS samples than the EDTA samples. These observations support and extend those of previous studies in which DBS DNA (both from Whatman FTA^®^ cards and other card types) has been profiled successfully using Infinium BeadArrays^[12, 32–36]^.

Two concerns cited when considering the use of archived DBSs are the effects of sample quantity and degradation with storage time^[14]^. Here we demonstrate that despite obtaining less DNA than recommended as input for the Infinium HumanMethylation450 BeadChip from 19 of the DBS samples, high quality methylation profiles were obtained. Moreover, we did not observe an effect of storage time on the success of methylation profiling; however, as the longest time that a sample was stored on a Whatman FTA^®^ card in the present study was 10 years, it remains possible that longer storage times might exert an effect. An earlier demonstration of successful methylation profiling of two DBS samples that had been stored for 26-28 years on standard filter paper^[34]^ does, however, provide additional support for the notion that extended storage time (as is the case for some potentially valuable archival sample collections) might not exert a deleterious effect.

Correlation and unsupervised hierarchical clustering analyses demonstrated a high degree of concordance between the DNA methylation profiles obtained from the Whatman FTA^®^ DBSs and the EDTA samples. These findings are consistent with those of Joo et al. (2013)^[35]^ and Hollegard et al. (2013)^[34]^ who showed methylation profiles obtained from DBS DNA samples to be highly correlated and/or cluster with those from paired buffy coat DNA or whole blood DNA samples from the same individual. These studies only considered small numbers of participants (n = two or four) for whom the DBSs had been stored for three years, rendering our findings in 62 participants with DBSs that had been stored for a mean period of eight years an important addition to previous studies.

EWASs of smoking and age in both the DBS and EDTA samples produced results that replicated previously established associations identified using peripheral blood DNA. The most significant locus identified in the EWASs of smoking status in both sample types was cg05575921 in AHRR, which was hypomethylated in smokers. This replicates multiple previous studies ^[43, 44]^, including meta-analyses^[39, 40]^, and cg05575921 has been suggested as a biomarker of smoking status^[41]^. Interestingly, one of the validation cohorts (Melbourne Collaborative Cohort Study^[42]^) assessed by Fasanelli et al. (2015)^[38]^ comprised 75% DBS samples. This cohort was used to successfully validate the two most significant DMPs (including cg05575921) identified in the discovery cohort.

The most significant locus in the EWASs of age in both the DBS and EDTA samples was cg16867657 in ELOVL2, where methylation was positively correlated with age, replicating previous studies^[43, 44]^. Methylation at this locus has been suggested as a biomarker of aging, with potential utility for forensic investigation^[45]^.

No significant associations were observed between DNA methylation and the interaction between smoking or age and storage method. This observation indicates that storing blood samples as DBSs at room temperature for approximately eight years is unlikely to affect the results of EWASs for traits that are robustly associated with variation in DNA methylation.

It is of interest to consider these findings in the context Dugué et al’s (2016)^[46]^ demonstration of lower reliability of DNA methylation profiling of DBSs using Infinium HumanMethylation450 BeadChips (compared to the moderate reliability of peripheral blood mononuclear cell DNA samples). This finding, which was based on the calculation of probe-wise intraclass correlation coefficients for technical replicates (ICC; the median of which was 0.20 for DBS technical replicates), suggests that, in general, there is a higher degree of measurement error when assaying DBS DNA. In light of this finding, Dugué et al. suggested that it may be advisable to only include probes with an ICC above a certain threshold when carrying out EWASs using DBS samples. It is clear from the present results, however, that measurement error was not so large as to obfuscate the pair-wise relationship between samples in the study or to confound EWASs of variables known to be robustly associated with variation in DNA methylation.

A disadvantage to storing blood samples using Whatman FTA^®^ cards is that it is not possible to sort the sample into separate blood cell types. As DNA methylation varies between blood cell types, the issue of cellular heterogeneity must be addressed in order to avoid confounding in EWASs using whole blood DNA^[25, 26]^. Fortunately, several algorithms have been developed previously to deal with the issue of cellular heterogeneity in EWAS studies (see Kaushal et al., 2017^[47]^ for a comparison of these methods) and our data supports the applicability of these to blood samples stored using Whatman FTA^®^ cards.

Taken together, the results of this study support the use of Whatman FTA^®^ cards as an alternative, more cost- and space-effective storage method for blood samples collected for DNA methylation studies. We demonstrate that DNA from DBSs stored for up to 10 years can be successfully profiled using Infinium BeadChips, producing methylation profiles that are highly correlated with matched EDTA samples. We envisage that this approach to collecting and storing blood samples for DNA methylation profiling will help circumvent many issues faced by researchers amassing large collections of blood samples, particularly where access to freezer facilities is limited. These findings are also relevant to the ongoing discussion regarding the potential use of new-born Guthrie cards for research purposes and supports previous work assessing the potential for methylomic profiling of these samples^[33–35, 48, 49]^.

## Data availability statement

Due to the confidential and potentially identifiable nature of the DNA methylation data and participant phenotypic information, it is not possible to publically share the individual level data used for these analyses. UK researchers and international collaborators can, however, request access to all GS data by contacting the GS Access Committee. GS operates a managed data access process including an online application form, which will be reviewed by the GS Access Committee. Summary information to help researchers assess the feasibility and statistical power of a proposed project can be requested by contacting. EWAS summary statistics will be made available on request.

## Acknowledgements

We are grateful to everyone who took part in GS: SFHS, the GPs and Scottish School of Primary Care for their help in recruitment, and the whole GS team, which includes academic researchers, clinic staff, laboratory technicians, clerical workers, IT staff, statisticians and research managers.

